# Recombinant biosensors for multiplex and super-resolution imaging of phosphoinositides

**DOI:** 10.1101/2023.10.18.562882

**Authors:** Hannes Maib, Petia Adarska, Robert Hunton, James Vines, David Strutt, Francesca Bottanelli, David H. Murray

## Abstract

Phosphoinositides are a small family of phospholipids, acting as signalling hubs and key regulators of cellular function. Detecting their subcellular distribution is crucial to gain insights into membrane organisation and is most commonly done by over-expression of biosensors. However, this leads to perturbations of phosphoinositide signalling and is challenging in systems that cannot be transfected. Here, we present a toolkit for the reliable, fast, multiplex, and super-resolution detection of all 8 phosphoinositides using a unifying staining approach for fixed cells and tissue, based on recombinant biosensors with self-labelling SNAP tags. These recombinant biosensors are highly specific, and reliably visualise the subcellular distributions of phosphoinositides across scales, ranging from 2D or 3D cell culture to *Drosophila* tissue. Using stimulated emission depletion (STED) microscopy, we reveal the nanoscale organisation of PI(3)P on endosomes and PI(4)P on the Golgi and confirm the preservation of subcellular membranes. Multiplex staining enables the investigation of phosphoinositide conversions and reveals an unexpected presence of residual PI(3,5)P_2_ positive membranes in swollen lysosomes following PIKfyve inhibition. This approach enables the versatile, high-resolution visualisation of multiple phosphoinositide species in an unprecedented manner.

## Introduction

Membrane identity is key to cellular function. A crucial hallmark of this identity are the phosphoinositides, a small family of phospholipids generated by phosphorylation of their phosphatidylinositol (PI) headgroup. Through the stepwise addition and removal of phosphate groups by lipid kinases and phosphatases, 8 distinct phosphoinositides can be generated that recruit specific effector proteins and thereby regulate a myriad of cellular functions. As such, the different phosphoinositides are involved in basically every cellular event involving membrane dynamics; ranging from cytokinesis to cell migration, autophagy, T-cell activation, membrane contact site regulation, as well as in their most well-known role as regulators of membrane trafficking (Edwards-Hicks et al., 2023; Gulluni et al., 2021; Jimenez et al., 2000; Mesmin et al., 2013; Tremel et al., 2021) see (Posor et al., 2022) for recent review). In line with their cell biological importance, continuing efforts are being undertaken to establish and refine tools to visualise their subcellular distribution.

Key to the visualisation of phosphoinositides is the identification of effector domains that specifically recognise the differentially phosphorylated headgroups (Hammond and Balla, 2015). These effector domains are often conserved Pleckstrin or Phox homology (PH and PX) domains but can also be structural folds such as GRAM, FYVE and Zinc finger domains, or the WD40 propeller found in WIPI1 and 2 (Dooley et al., 2014). To further complicate matters, some proteins bind to specific phosphoinositides through polybasic regions without any conserved secondary structure, such as Exoc7 (Exo70) as part of the exocyst complex (He et al., 2007; Maib and Murray, 2022). Through the combined effort of the community, these effector domains have been optimised to generate biosensors to detect phosphoinositides (Wills et al., 2018). With the recent addition of a PX domain from *Dictyostelium discoideum* that recognises PI(3,5)P_2_, (Vines et al., 2023), we now possess biosensors for each of the eight different phosphoinositides (albeit with uncertainties about PI(5)P). A common feature of many of these biosensors is the use of tandem and triple repeats to increase binding through an avidity effect, as is the case for the 3xPH domain of TAPP1, the 3xPHD domain from ING2, the 2xFYVE domain of Hrs and the 2xPX domain of SnxA to recognise PI(3,4)P_2_, PI(5)P, PI(3)P and PI(3,5)P_2_ respectively.

The most common way to visualise the subcellular localisation of phosphoinositides is to ectopically overexpress the lipid effectors fused to fluorescent proteins as biosensors (Wills et al., 2018). This approach allows for detection of membrane identity changes in live cells and has been of immense value over the last decades. However, it also suffers from a major drawback: as these biosensors are generated to have a high affinity, they can all potentially out-compete endogenous effector proteins and thereby perturb the very pathway that is under investigation. In contrast, biosensors with lower affinity and expression levels are themselves out-competed by endogenous effectors. Furthermore, this over-expression based approach is limited by the ability to deliver DNA into cells or requires time and labour extensive genomic engineering. It is therefore still a challenge for experimentalists to visualise these critically important lipids, especially in more complex systems such as 3D cell cultures or whole tissues.

One way to circumvent these pitfalls has been to stain for phosphoinositides after fixation and permeabilization using immunocytochemical techniques based on recombinant biosensors (Gillooly et al., 2000; Watt et al., 2002) and antibodies (Hammond et al., 2009; Maekawa and Fairn, 2014; Marat et al., 2017). While this approach is unable to provide dynamic information, it does avoid perturbation of phosphoinositide signalling and does not require overexpression. However, the need of fixatives and detergents requires careful optimisations to avoid artifacts and the available antibodies require different staining protocols for the visualisation of distinct pools of their respective targets (Hammond et al., 2009). As such, there is currently no reliable tool to comprehensively visualise all subcellular phosphoinositide pools with a unifying staining approach.

To address these shortcomings, we have developed recombinant biosensors against all eight phosphoinositides in combination with self-labelling protein tags (SNAP). With the exception of the only known biosensor against PI(5)P, all of these probes are easy to purify using standard bacterial expression systems and demonstrate excellent specificities, as determined using an *in vitro* supported lipid bilayer approach. Using a single, unified staining protocol, we show that these probes reliably visualise all phosphoinositide species after fixation and permeabilisation and reproduce their known subcellular localisations. The ease of conjugating distinct fluorophores to the SNAP tag enables the use of inorganic dyes, including those that are compatible with super resolution STED microscopy. This reveals the nanoscale organisation of PI(3)P at endosomes and of PI(4)P at the Golgi while furthermore verifying the preservation of subcellular membranes. Labelling with distinct fluorescent dyes *in vitro* enables multiplex imaging and allows for the detection of phosphoinositides across scales and model systems as shown by staining in HeLa cells, NMuMG spheroids and *Drosophila* pupal wings. To highlight the versatility of this approach, we investigate the interdependence of phosphoinositide conversion by multiplex staining for PI(3)P, PI(4)P and PI(3,5)P_2_ after inhibition of lipid kinases in MIA-Paca2 cells. These experiments unveil an unexpected sequestration of residual PI(3,5)P_2_ positive membranes in swollen lysosomes following PIKfyve inhibition.

## Results

### Recombinant biosensors are straightforward to produce and are highly phosphoinositide specific

The cell biology community has spent considerable effort to identify effector domains of proteins that recognise specific phosphoinositides for the use as biosensors (see table in Fig. 1). To test their suitability as recombinant probes, we designed effectors with an N-Terminal 6xHis tag followed by a SNAP tag and a flexible linker region for bacterial expression. With the exception of the 3xPHD domain of ING2 (see Discussion), these recombinant biosensors are readily purified with high yields from *E. coli* using a standard three-step purification procedure of Ni^2+^ affinity followed by ion exchange and size-exclusion chromatography. The inclusion of a SNAP tag then allows for versatile *in vitro* labelling of the probes using inorganic fluorophores for multiplexed and/or super resolution microscopy (Fig. 1).

**Figure 1:**
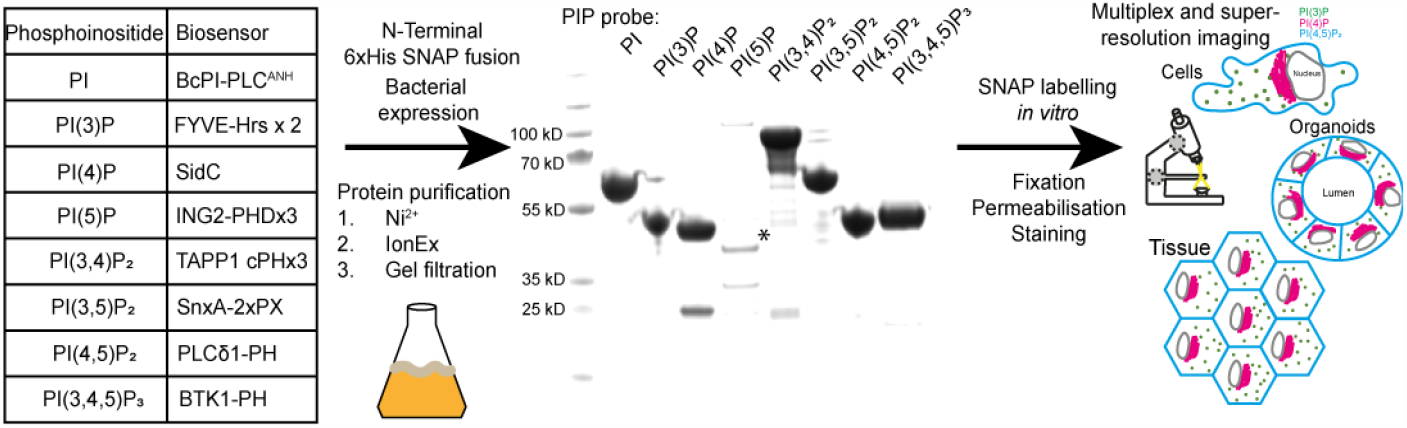
Scheme for the generation of recombinant biosensors for detection of phosphoinositides in fixed and permeabilised cells. Left: Table of Biosensors used to for the detection of each phosphoinositide. Middle: Coomassie stained gel of purified probes after three step purification from *E. Coli*. The asterisk denotes the soluble fraction of the PI(5)P biosensor, which is only stable at low concentrations.

To test the specificities towards their respective targets, we labelled the biosensors with SNAP-Surface Alexa Fluor 488 *in vitro* and formed supported lipid bilayers on silica beads with liposomes composed of 95% 1-palmitoyl-2-oleoyl-glycero-3-phosphocholine (POPC), and 5% of each phosphoinositide. These membrane coated beads are an ideal substrate to evaluate binding specificities, as they faithfully mimic the charge and fluidity of biological membranes (Pucadyil and Schmid, 2010). The quality of the lipid bilayers is verified by including 0.1% of fluorescently labelled lipids (here Atto647N-DOPE) and using this signal to generate a membrane mask to quantify the recruitment of the biosensors (Fig.2).

The biosensor for PI is a modified version of a bacterial, PI-specific PLC (Pemberton et al., 2020) and shows good specificity towards PI albeit with lower affinity compared to the other probes (Fig. 2a). The biosensor for PI(3)P is the well known tandem repeat of the FYVE domain from HRS (Burd and Emr, 1998) and shows excellent specificity and affinity for PI(3)P with some weak binding towards PI(3,5)P_2_ (Fig. 2b). The probe for PI(4)P is the effector domain of SidC (also known as P4C) (Dolinsky et al., 2014) and shows strong binding to PI(4)P with some background binding to the other mono-phosphates PI(3)P and PI(5)P (Fig. 2c). The probe for PI(5)P is the triple repeat of the PHD domain from ING2 (Gozani et al., 2003) and shows little specificity. Even though this probe shows stronger binding to PI(5)P compared to PI, PI(3)P, PI(3,4)P_2_ and PI(4)P, it binds even more strongly to the other 5’ containing di- and tri-phosphates, suggesting a charge based binding effect in addition to 5’ binding (Fig. 2d). Therefore, even though this biosensor displayed promising results from overexpression in cells, it is unlikely to be suitable as recombinant biosensor. The biosensor for PI(3,4)P_2_ is the triple repeat of the C-terminal PH domain from TAPP1 (Goulden et al., 2019) and shows good specificity and affinity with some binding to the di- and tri-phosphates (Fig. 2e), likely due to the strong positive charge of this probe (with a pI of 9.7). The biosensor for PI(3,5)P_2_ is the recently described tandem repeat of a PX domain from *Dictyostelium discoideum* (Vines et al., 2023) and shows excellent specificity and affinity with almost undetectable binding to any of the other phosphoinositides (Fig. 2f). The biosensor for PI(4,5)P_2_ is the well-established PH domain from PLC-δ1 (Garcia et al., 1995) and shows reassuring specificity (Fig 2g). And finally the biosensor for PI(3,4,5)P_3_ is the PH domain from BTK (Fukuda et al., 1996; Salim et al., 1996) and shows good specificity with some minor binding to PI(3,4)P_2_ (Fig 2h). These comprehensive and directly comparable *in vitro* data validate the specificity for each of the recombinant biosensors, and suggest their possibility for use in staining approaches.

**Figure 2:**
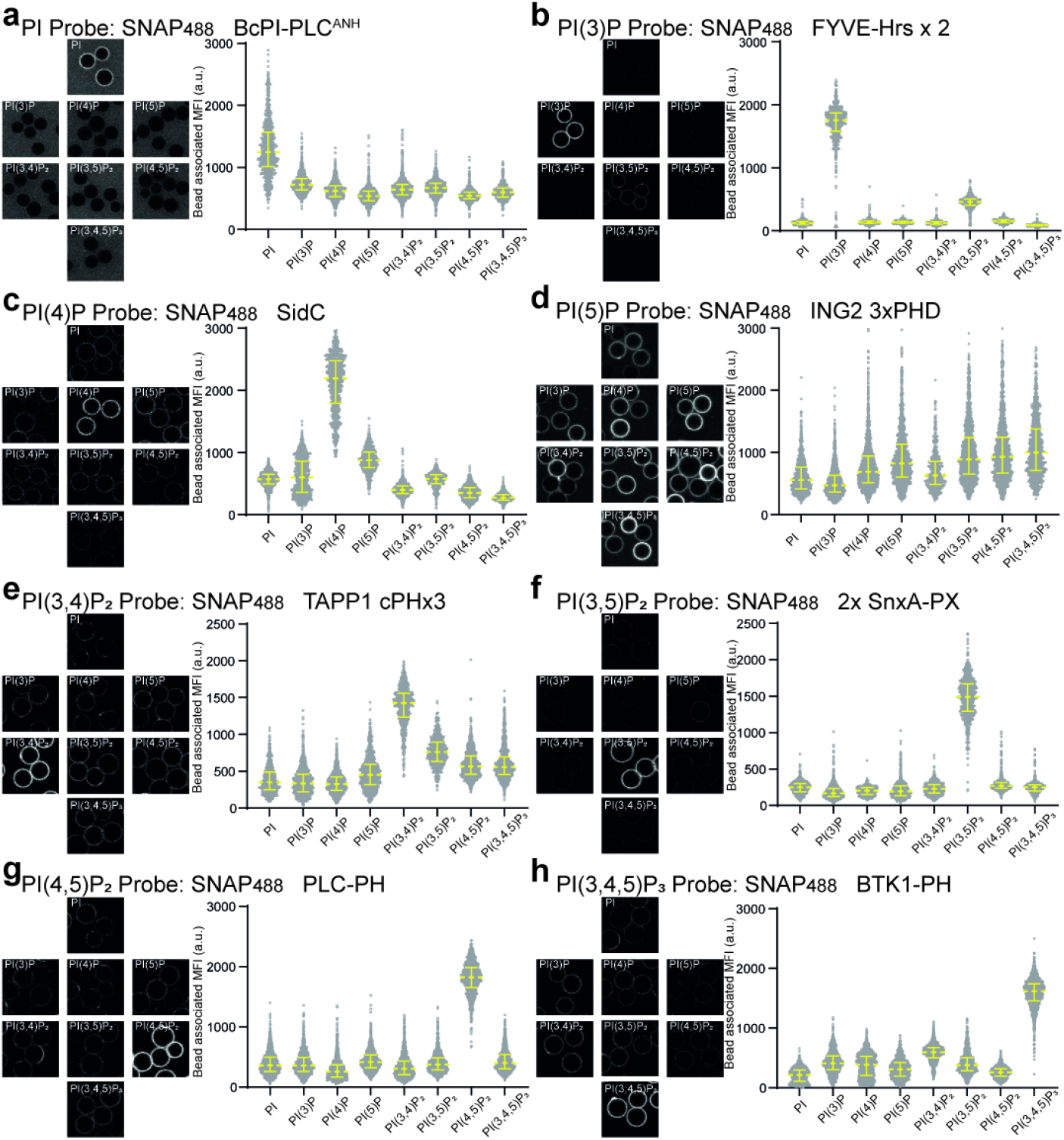
*In vitro* verification of recombinant biosensors. (a-h) the indicated biosensors were labelled with Alexa488 and added at 200nM final concentration to membrane coated beads containing 5% of the indicated phosphoinositide. Mean fluorescence intensity (MFI) of biosensor bound to each individual bead was determined by semi-automated segmentation using the fluorescent DOPE lipid as mask in ImageJ. Data is presented as median with interquartile range from n= 473-1642 individual beads for each phosphoinositide and each biosensor.

### Subcellular visualisation of all phosphoinositide species by a unified staining approach

Visualisation of phosphoinositides using these recombinant biosensors requires cell fixation and permeabilisation. Importantly, multiplex imaging requires a unifying staining protocol that is easy to use while preserving all of the distinct phosphoinositide pools, whether they are at the plasma membrane, Golgi, or endosomes. To achieve this objective, HeLa cells were fixed with prewarmed (37°C) 4% PFA with 0.2% Glutaraldehyde and permeabilised using 0.5% Saponin on ice and stained with 500nM of each of the separate biosensors labelled with Alexa 488 (Fig. 3) (see methods for more details).

**Figure 3:**
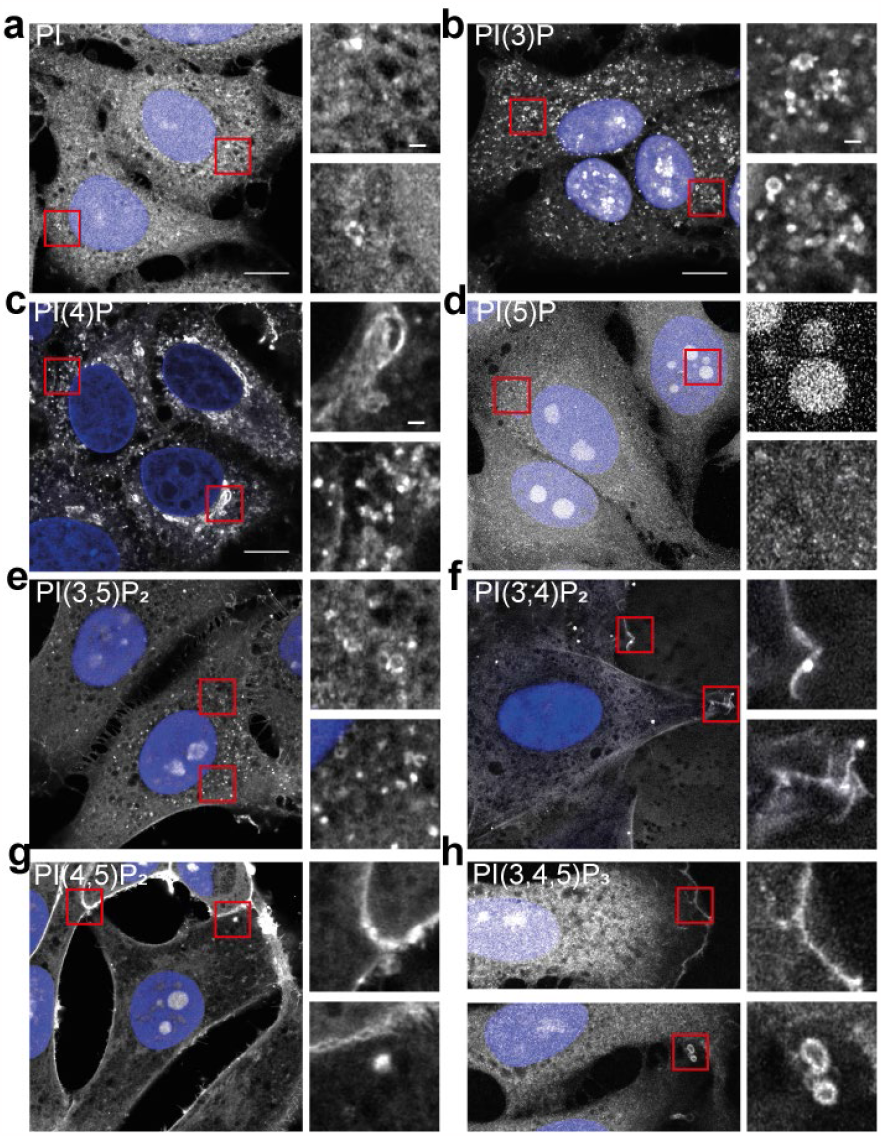
Cellular stainings of phosphoinositides in fixed and permeabilised cells using recombinant biosensors. a-h) HeLa cells were grown on cover slips and fixed and permeabilised as described in the methods section. Phosphoinositides were detected using recombinant biosensors conjugated to Alexa488 at 500nM final concentration and imaged using Airyscan microscopy. Scale bar = 10μm and 1μm in inserts.

The staining for PI in HeLa cells shows a rather diffuse pattern with some membrane staining but poor contrast (Fig. 3a). This is likely due to the relatively low affinity of this probe and the almost ubiquitous presence of this phosphoinositide on subcellular membranes. The staining for PI(3)P on the other hand shows excellent signal to noise ratio with clear labelling of endo/lysosomal structures throughout the cell (Fig. 3b). The PI(4)P staining shows excellent labelling of all known pools of this phosphoinositide, including the Golgi, trafficking vesicles, and the plasma membrane at junctions between cells (Fig. 3c). The cellular staining using the recombinant biosensor against PI(5)P shows no localisation to membranes, with the only clear signal in the nucleolus (Fig. 3d). This is most likely due to the strong negative charge of open DNA and agrees with the mainly unspecific, charge based binding seen with supported lipid bilayers (Fig. 2d). Staining with the PI(3,5)P_2_ biosensors reveals faint vesicular staining, in agreement with the low steady state levels of this rare phosphoinositide in HeLa cells (Fig. 3e). In agreement with its known localisation the PI(3,4)P_2_ biosensor stains weakly the plasma

### Recombinant biosensors in combination with STED microscopy reveal the nanoscale organisation of phosphoinositides

Understanding the nanoscale organisation of phosphoinositides at sub-cellular membranes is key towards gaining novel insights into their functions. To investigate this organisation, and to verify the preservation of sub-cellular membranes, we harnessed super-resolution microscopy. The inclusion of self-labelling protein tags allows for the use of bright, photostable, inorganic Janelia^®^Flour dyes (Grimm et al., 2020; Grimm et al., 2021) that are exceptionally well-suited for stimulated emission depletion (STED) microscopy. Staining HeLa cells with the PI(3)P biosensor conjugated to JFX650 shows the well-established endosomal localisation of this phosphoinositide in “microdomains” (Gillooly et al., 2003). When viewed using STED microscopy, however, it is revealed that these domains are indeed nanoscale membrane tubulations as well as smaller vesicles that cluster around these larger endosomes and potentially fuse with, or bud from them (Fig. 4a), as has been seen previously using correlative light and electron microscopy (Pucadyil and Schmid, 2010).

**Figure 4:**
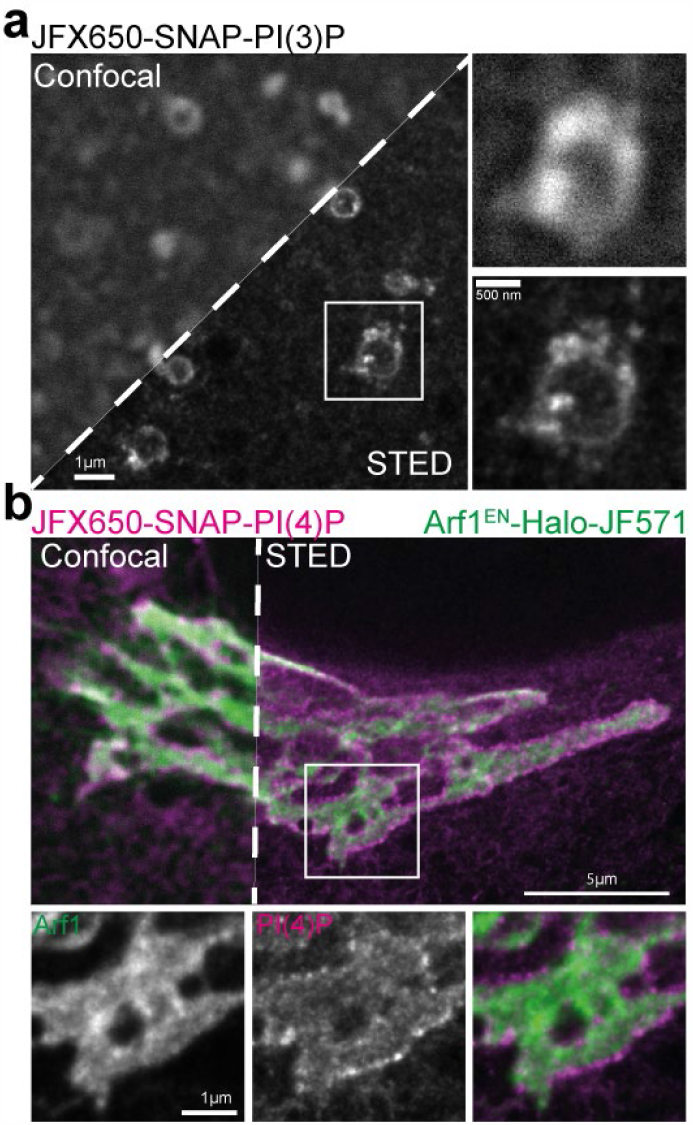
Super-resolution STED imaging of nanoscale phosphoinositide organisation. a) Endosome staining using biosensor against PI(3)P conjugated with JFX650 as seen by confocal and STED microscopy. b) Two colour STED microscopy of Golgi with endogenously tagged Arf1^EN^-Halo stained with JF571 and PI(4)P using recombinant biosensor conjugated to JFX650. membrane with enrichment at membrane ruffles (Fig. 3f). The PI(4,5)P_2_ biosensors reveals strong plasma membrane and a weak intracellular staining, in agreement with the well described presence of minor pools of this phosphoinositide on intracellular membranes (Fig. 3g). Finally, the biosensor for PI(3,4,5)P_3_ shows selective staining of sub-regions of the plasma membrane such as the leading edge and structures reminiscent of macropinocytic cups (Fig. 3h). In summary, these stains demonstrate a unifying protocol for staining and visualization of all phosphoinositide species.

To gain more insights into the nanoscale organisation of phosphoinositides, we investigated the distribution of PI(4)P at the Golgi using the same, universal staining protocol. HeLa cells expressing endogenously tagged Arf1^EN-^Halo were first stained live against the Halo tag using the JF571 fluorophore before being fixed, permeabilised and stained against PI(4)P using the recombinant biosensor conjugated to JFX650. Arf1 is a crucial factor in the generation of PI(4)P at the Golgi through recruitment of PI4K IIIβ (Highland and Fromme, 2021) and this combination of dyes allows for two-colour STED imaging (Wong-Dilworth et al., 2023). Excitingly, the staining for PI(4)P reveals distinct nanoclusters of this lipid at the curved rims of the cisternae where budding of trafficking intermediates is thought to occur as well as diffuse staining throughout the Golgi (Fig. 4b). The localization of the PI(4)P sensor is in fact reminiscent of COPI clusters (Wong-Dilworth et al., 2023). Thereby, this robust approach for the visualisation of phosphoinositides enables super-resolution interrogation of membrane organisation while preserving the structure of sub-cellular membranes.

### Multiplex detection of phosphoinositides across scales and model systems

A staining protocol, which preserves the subcellular organisation of membranes and is compatible with the detection of each of the distinct phosphoinositides, opens up the possibility for multiplex imaging. As proof of principle, HeLa cells were stained with recombinant biosensors against PI(3)P, PI(4)P and PI(4,5)P_2_ conjugated to Alexa488, 546 or 647 and imaged using Airyscan microscopy (Fig. 5a). Amongst numerous observations, we noted PI(4)P positive tubules in direct contact with PI(3)P positive endosomes in the vicinity of the PI(4,5)P_2_ positive plasma membrane (Fig. 5a (insert)). This multiplex approach highlights the intricate interactions of membranes with distinct phosphoinositide identities, critically important to the organisation of organelle contact sites (Posor et al., 2022).

**Figure 5:**
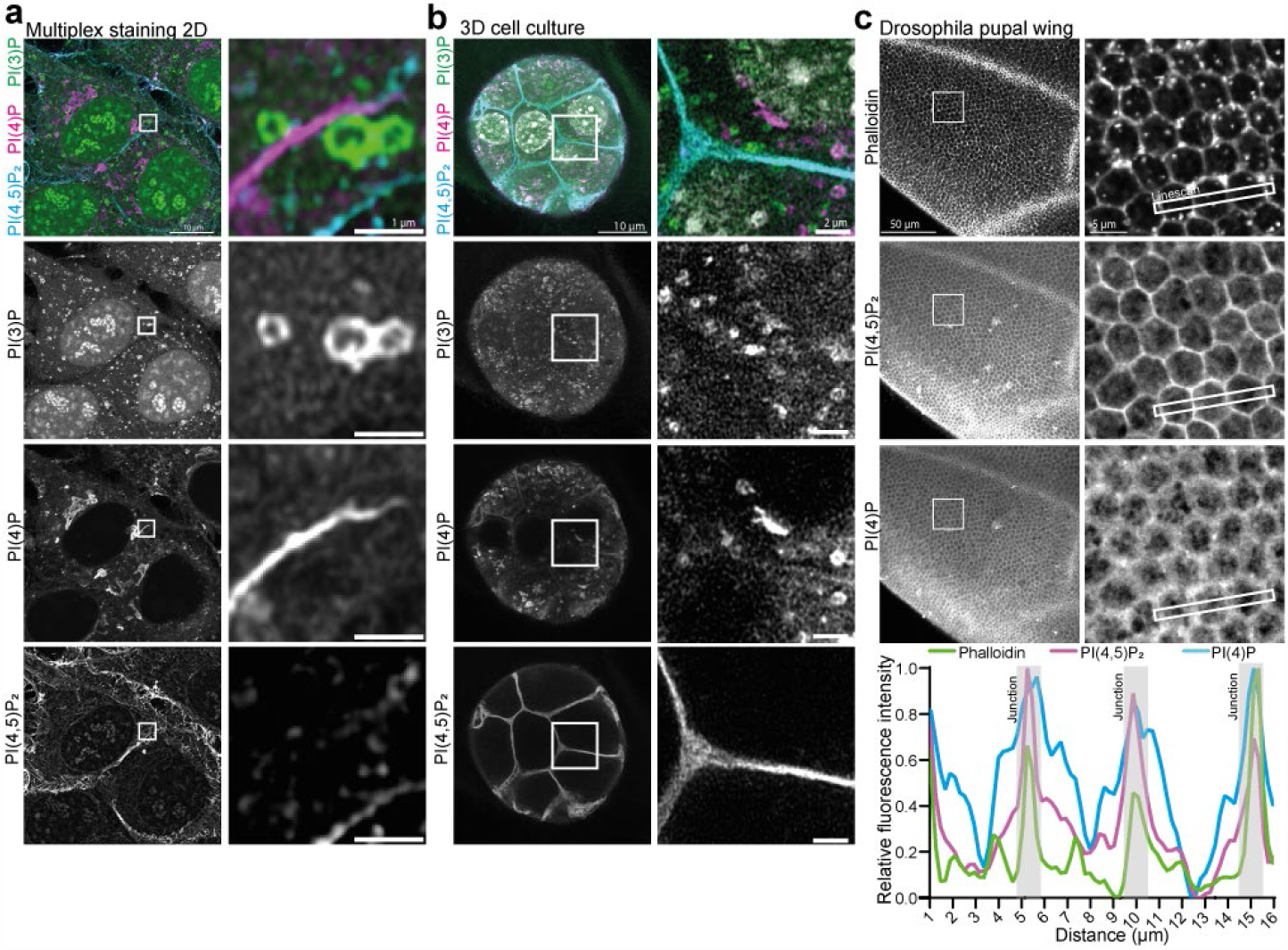
Multiplex staining of phosphoinositides across scales. a) HeLa cells were fixed, permeabilised and stained using recombinant biosensors against PI(3)P, PI(4)P and PI(4,5)P_2_, conjugated respectively to Alexa488, 546 and 647. b) NMuMG spheroids were grow in Matrigel and stained with the same combination. c) *Drosophila* pupal wings were dissected and stained with the PI(4,5)P_2_, and PI(4)P biosensors conjugated to Alexa647 and 546, together with Phalloidin conjugated to Alexa488 to visualise the actin cytoskeleton and cellular junctions.

An important advantage of recombinant biosensors is that it avoids the need of overexpression. As such, it allows for the detection of phosphoinositides in cells and tissues that cannot easily be transfected and negates the need of genome engineering. To highlight this advantage, we grew NMuMG spheroids in Matrigel for 5 days and used the same staining protocol as previously to detect PI(3)P, PI(4)P and PI(4,5)P_2_ (Fig. 5b). As in 2D cell culture, these three phosphoinositides show an intricate interaction at the subcellular level with strong labelling of PI(4,5)P_2_ at cell-cell junctions. To further test the utility of this approach in whole tissues, we stained *Drosophila* pupal wings against PI(4)P and PI(4,5)P_2_ in combination with conventional Phalloidin to visualise the actin cytoskeleton and cellular junctions. Consistent with the results from 3D culture, the recombinant biosensor against PI(4,5)P_2_ shows strong labelling of cellular junctions throughout the whole tissue. Noticeably, PI(4)P is also enriched at these junctions albeit with a more diffuse staining pattern, indicating the presence of intracellular PI(4)P membranes in close proximity to the plasma membrane (Fig. 5c). These observations are in good agreement with the dynamic regulation of these two lipids at cell junctions in *Drosophila* tissue to drive recruitment of polarity proteins through electrostatic interactions (Dong et al., 2015; Lu et al., 2022).

These experiments demonstrate conclusively that this toolkit enables the visualisation of several phosphoinositide species simultaneously without the adverse effects of overexpression. This will enable the visualisation of these crucial lipids across scales and model systems, from 2D to 3D cell culture and thin tissues.

### Staining reveals hidden pools of PI(3,5)P_2_ following PIKfyve inhibition

Multiplex staining of phosphoinositides enables investigation of conversion cascades in a straight-forward manner and avoids overexpression of multiple biosensors. To highlight this advantage, and to make use of a novel PI(3,5)P_2_ probe (Vines et al., 2023), we decided to focus on the conversion cascade leading to the generation of this rare phosphoinositide. Staining for PI(4)P, PI(3)P and PI(3,5)P_2_ in combination with Airyscan microscopy reveals the close relationship of these three phosphoinositide (Fig. 6a). While PI(4)P membranes are often in close proximity to PI(3)P containing endosomes, it is rarely found on the same vesicle. PI(3,5)P_2_ on contrast is located on the same membrane as PI(3)P and enriched in domains that likely represent membrane tubulations and budding/fusion of smaller vesicles (Fig. 6a). To investigate the conversion cascade of PI → PI(3)P → PI(3,5)P_2_, we used specific kinase inhibitors to block each conversion step in MIA–Paca2 cells, due to a low steady state level of PI(3,5)P_2_ in HeLa cells. As expected, treatment with the potent pan PI3K inhibitor Wortmannin (at 300nM for 60 min) eliminates vesicular staining for PI(3)P as well as for PI(3,5)P_2,_ with minor effects on PI(4)P distribution (Fig. 6b middle). Treatment with Apilimod, a highly selective and effective PIKfyve inhibitor (Cai et al., 2013) (at 1μM for 60 min), leads to the accumulation of PI(3)P on endosomes as well as the formation of swollen lysosomes, and the depletion of PI(3,5)P_2_. We also observed some morphological changes in the organisation of PI(4)P membranes (Fig. 6b bottom). This confirms the specificity of the staining approach and that the generation of PI(3,5)P_2_ is strictly dependent on the prior formation of PI(3)P, which accumulates on endosomes when its conversion into PI(3,5)P_2_ is blocked.

**Figure 6:**
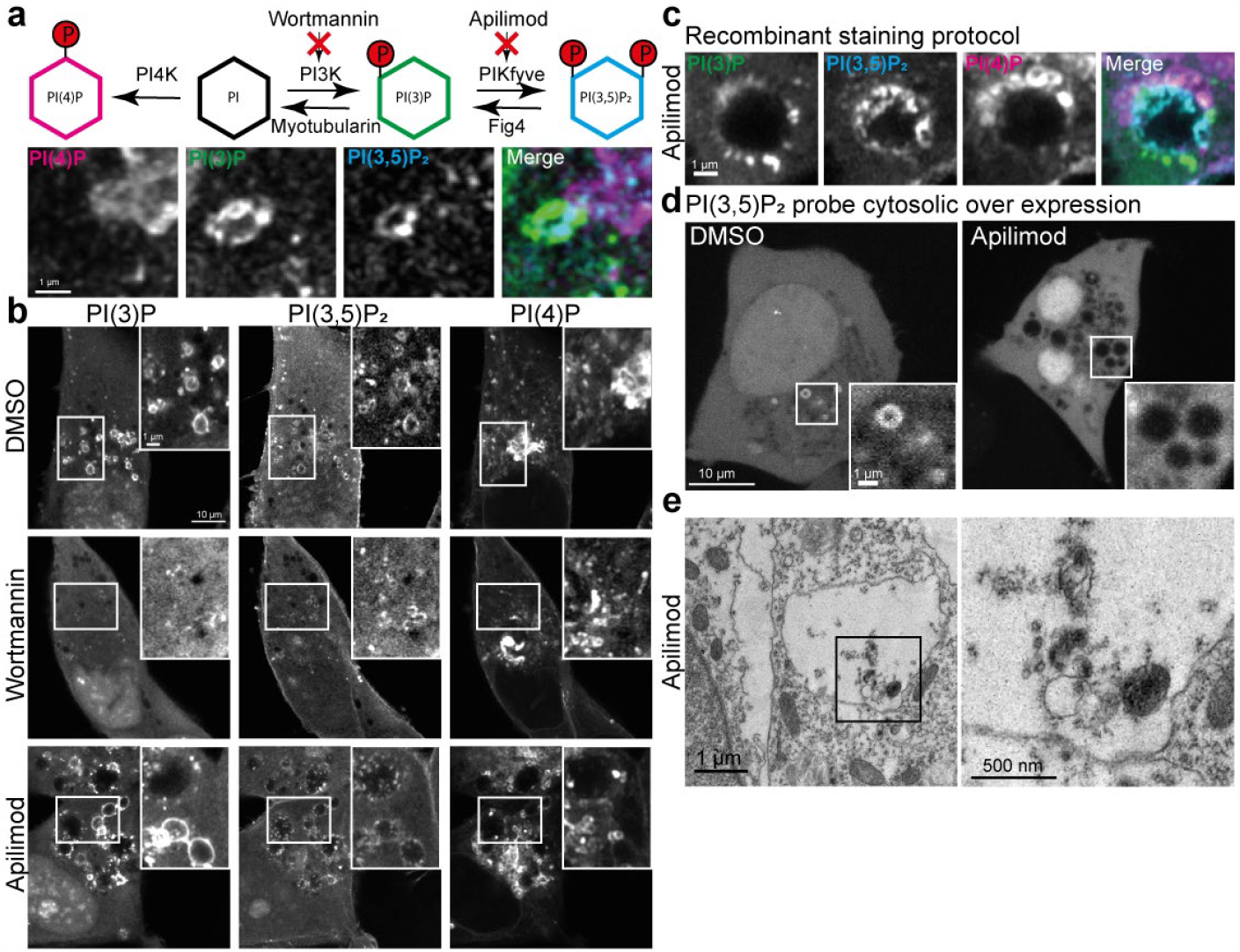
Phosphoinositide interconversion and hidden PI(3,5)P_2_ positive intraluminal membranes following PIKfyve inhibition. a) Interconversion scheme and multiplex staining of PI(4)P, PI(3)P and PI(3,5)P_2._ b) MIA-Paca2 cells were treated with DMSO, 300nM Wortmaninn or 1uM Apilimod for 60min before fixation, permeabilisation and multiplex staining. c) Example of swollen lysosomes following Apilimod treatment and stained against PI(4)P, PI(3)P and PI(3,5)P_2_. d) PI(3,5)P_2_ biosensor, fused to eGFP, was cytosolically over-expressed and cells were imaged life after Apilimod treatment. e) Thin section electron micrograph of swollen lysosomes in MIA-Paca2 cells after treatment with Apilimod.

Upon closer inspection of the images, we noticed that the PI(3,5)P_2_ biosensor detects a distinct signal in the lumen of the swollen lysosomes upon PIKfyve inhibition (Fig. 6c). Importantly, these intraluminal membranes are inaccessible to a cytosolically expressed PI(3,5)P_2_ biosensor - highlighting an additional limitation of this conventional approach (Fig. 6d). To address the nature of this staining, we investigated these swollen lysosomes using standard thin section electron microscopy. This orthogonal technique confirms the presence of these intraluminal membranes, corresponding to the staining observed using the recombinant PI(3,5)P_2_ biosensor. This unexpected detection of residual PI(3,5)P_2_ membranes following PIKfyve inhibition highlights another advantage of the staining approach by making intraluminal membranes accessible that are otherwise hidden from the cytosolic overexpression of these probes.

## Discussion

The subcellular detection of phosphoinositides using fluorescence microscopy has been instrumental for the investigation of these signalling lipids. Through the combined efforts of the community, specific effector domains have been identified to detect each phosphoinositide, and widely used in over-expression based approaches. This approach, however, often perturbs phosphoinositide signalling, is not possible in cell types that cannot be transfected and is perilous for multiplex detection due to the need to express multiple probes. To address these shortcomings, we have generated a toolkit of recombinant biosensors for the detection of all eight phosphoinositides for multiplex and super resolution imaging in fixed and permeabilised cells.

These recombinant biosensors are easily purified by standard means, show excellent specificities *in vitro*, and the use of self-labelling protein tags (SNAP) allows for high versatility in the choice of inorganic fluorophores. These fluorophores benefit from a vastly improved brightness and photostability compared to fluorescent proteins and allow for the detection of minor phosphoinositide pools. Importantly, the use of a unifying fixation and permeabilisation protocol for the detection of distinct subcellular pools allows for multiplex staining. This protocol is adapted from Hammond et al., 2009 where it was used for immunocytochemical detection of PI(4)P and PI(4,5)P_2_ using commercial antibodies. However, these antibodies detect different phosphoinositide pools depending on the staining protocol. As such, after permeabilisation with Saponin, they detect PI(4)P at the plasma membrane but not at the Golgi. Conversely, permeabilisation using Digitonin enabled visualisation of PI(4)P at the Golgi but abolishes staining of PI(4)P and PI(4,5)P_2_ at the plasma membrane (Hammond et al., 2009). Importantly, the recombinant biosensors that we have presented here, allow visualisation of all of these distinct phosphoinositide pools with the same, Saponin based, staining protocol. Still, to reach all subcellular compartments, their surrounding membranes have to be disrupted using detergents, which might introduce artifacts. To address this concern, we used super resolution STED microscopy and could confirm that sub-cellular organelles retain their structure following this staining protocol. Therefore, these recombinant biosensors offer a clear advance for the subcellular detection of phosphoinositides. However, some minor pools might always escape detection due to being occupied by endogenous effectors.

Out this toolkit of biosensors, the probe to detect PI(5)P is the only one that is challenging to produce recombinantly. The only established probe to detect this elusive phosphoinositide is based on the Zinc finger domain of ING2 (Gozani et al., 2003) and has shown promising results in over-expression based approaches (Pendaries et al., 2006; Vicinanza et al., 2015). However, it is highly unstable as recombinant protein, requires supplementation of Zn^2+^ and aggregates at high concentrations; although it can be separated from its aggregates in low concentrations using size-exclusion chromatography. Disappointingly though, even after careful biochemical qualification, the recombinant probe shows little specificity towards PI(5)P *in vitro* and only shows unspecific staining of the nucleolus when used to stain cells. Therefore, further work will have to be carried out in order to identify a reliable probe that works as recombinant biosensor to detect PI(5)P.

The nanoscale organisation of phosphoinositides is crucial for the recruitment of effector proteins and the downstream regulation of cellular function. To shed more light on this we made use of the development of inorganic fluorophores in combination with super resolution STED microscopy and stained cells with recombinant biosensors against PI(3)P and PI(4)P. The organisation of PI(3)P in microdomains on endo/lysosomes has been well described for over two decades (Gillooly et al., 2003) however their precise organisation has been somewhat unclear. By using STED, we could show that these domains are indeed nano tubulations and smaller vesicles in agreement with correlative light and electron microscopy of small GTPases on early endosomes (Franke et al., 2019). This nanoscale organisation is supported by the role of PI(3)P as effector for EEA1 in the tethering of early endosomes (Murray et al., 2016), as well as in recruitment of ESCRT-0 in the formation of multi vesicular bodies (Banjade et al., 2022) and in regulation of the CCC complex in WASH mediated recycling (Singla et al., 2019). Similarly, PI(4)P is a crucial identity determinant of the Golgi and regulates secretion (Waugh, 2019). The nanoclusters we have observed here are consistent with the clustering of this lipid through multiple effectors in the formation of AP1 and COPI coats ((Lorente-Rodriguez and Barlowe, 2011) (Tan and Brill, 2014)) and opens up new avenues for future research. We believe that this approach of staining for phosphoinositides in combination with STED microscopy will reveal further insights into the nanoscale organisation of phosphoinositides in the years to come.

The ability to reliably stain for phosphoinositides enables the investigation of these crucial lipids across scales and model systems. In addition to super resolution microscopy, the versatility of self-labelling protein tags makes differential labelling and multiplex imaging possible. As proof of principle, we have performed triple staining of the most abundant phosphoinositides: PI(3)P, PI(4)P and PI(4,5)P_2_ in 2D as well as 3D cell culture. This highlights the close interactions these membranes, as has been shown for the role of PI(4)P containing membranes in regulating the scission of PI(3)P positive endosomes (Boutry et al., 2023; Gong et al., 2021). To further highlight the versatility of our approach, we stained *Drosophila* pupal wing tissue against PI(4,5)P_2_ and PI(4)P in combination with Phalloidin to visualise actin filaments. This approach clearly detects high levels of both PI(4)P and PI(4,5)P_2_ at cellular junctions throughout the whole tissue. We noticed that the levels of PI(4)P at junctions seemed to increase from 2D to 3D cell culture and tissue. While warranting further research, this increase in negatively charged phospholipids in polarised systems is in good agreement with the recruitment of polarity proteins through electrostatic interactions (Dong et al., 2015; Lu et al., 2022). Importantly, the ability to visualise phosphoinositides in model systems that are difficult to genetically modify has the potential to reveal novel insights into their functions on the tissue scale and furthermore opens up the possibility to investigate these conserved lipids in eukaryotes across the evolutionary tree.

Multiplex staining also enables the investigation of phosphoinositide conversion cascades, as we have shown for the generation of PI(3,5)P_2_ following inhibition of different lipid kinases. Unexpectedly, this also revealed the presence of residual PI(3,5)P_2_ positive membranes in the lumen of swollen lysosomes following PIKfyve inhibition using Apilimod. While the formation of these swollen lysosomes is well known (Cai et al., 2013), the nature of these intraluminal vesicles is less clear. Importantly, these intraluminal vesicles are inaccessible to cytosolic kinases and phosphatases. Thus, since PIKfyve is the only known kinase that generates PI(3,5)P_2_, these pools must be present prior to Apilimod treatment and sequestered into these terminal lysosomes where they cannot be broken down. Further, PIKfyve forms a constitutive complex with the scaffolding protein Vac14 and the lipid phosphatase Fab1 that catalyses 5’ dephosphorylation of PI(3,5)P_2_ back to PI(3)P (Lees et al., 2020). This indicates an intricate cross regulation of the PIKfyve:Vac14:Fab1 complex to avoid futile phosphorylation cycles. In this context, it is tempting to speculate that the inhibition of PIKfyve might also indirectly influence the activity of Fab1. Further work into this regulation is underway and the ability to stain for phosphoinositides and visualise intraluminal membranes will be crucial towards its progress.

Taken together, we believe that this toolkit is a valuable addition to the repertoire of cell biologists to investigate phosphoinositide biology. Finally, this approach has the potential to reveal novel insights related to health and disease, especially in cell types and models that cannot be transfected, such as patient derived primary cells and tissue sections.

## Materials and Methods

### Constructs and Cloning

DNA for all biosensors was optimized for bacterial expression, synthesised, and cloned into standard ITPG inducible bacterial expression vectors containing Kanamycin resistance. Following the N-Terminal 6xHis tag an HRV3C cleavage site has been added upstream of the SNAP-tag for optional removal of the His tag. All constructs are available through Addgene.

### Purification and labelling of recombinant biosensors

All recombinant biosensors were expressed in BL21 bacterial cells using standard approaches. Transfected bacteria were grown in LB containing 1.75 w/vol % lactose and antibiotic (+5mM Zinc Chloride for purification of ING2-3xPHD) at 37C to OD600=0.8, whereupon temperature was lowered to 18C for 10-12 h. Cells were pelleted, resuspended in standard buffer (20mM HEPES, 250mM NaCl, 0.5 mM TCEP, +1mM Zinc Chloride for purification of ING2-3xPHD), lysed, and clarified by centrifugation at 55.000G for 1h at 4C. Cleared lysates were passed through a 0.45μM filter (Sartorius) and protein was purified by Ni^2+^ affinity chromatography using 5ml His-Trap HP column (Cytiva) followed by anion exchange on a 5ml Capto-Q or Capto-S column (Cytiva) depending on the pI of the biosensor. Peak fractions were pooled and purified by size exclusion chromatography using a Superdex 200 16/60 pg column (Cytiva). All proteins were aliquoted, frozen in liquid nitrogen and stored at -80C. Aliquots were labelled with SNAP-Surface Alexa Fluor 488, 546 or 647 (NEB) or JF650 (Janelia Fluor) in a 2:1 molar excess for 2-3h on ice in standard buffer. Excess dye was removed by dialysis against standard buffer at 4C overnight. Note that removal of excess dye improves the signal to noise of cellular staining but is not a strict necessity as it gets washed out during the staining process.

### *In vitro* lipid binding experiments

1 mg liposomes, containing each of the eight different phosphoinositides were produced by mixing 95 mol % 1-palmitoyl-2-oleoyl-sn-glycero-3-phosphocholine with 5 mol % of the respective phosphoinositides together with 0.1% Atto647N-DOPE. The mixtures were evaporated under nitrogen and dried overnight in a vacuum extruder. Dried lipids were resuspended in 1 ml buffer containing 150mM NaCl, 20mM Hepes, and 0.5mM TCEP and subjected to six freeze–thaws in liquid nitrogen. Liposomes of ∼100nm diameter were generated by passing the lipid mixture 11 times through a 100-nm filter (Whatman Nuclepore). Liposomes were aliquoted, snap-frozen, and stored at −20°C.

Membrane-coated beads were generated by adding 10 μg of liposomes to ∼0.5 × 10^6^ 10μm silica beads (Whitehouse Scientific) in 100μl of 200mM NaCl for 30 min rotation at room temperature. Beads were washed twice and resuspended in a buffer containing 150mM NaCl, 20mM Hepes, and 0.5mM TCEP and blocked with 200 ug/mL beta casein. Drops of 10μl beads were added into the corners of uncoated μ-Slide 8-well chambers (Ibidi) and 10μl of the purified biosensors were added to 200nM final concentration. Lipid-binding kinetics were allowed to equilibrate for 30 min at room temperature before imaging close to the equator of the beads.

Confocal images were acquired using a Leica SP8 Confocal Microscope with a Leica HC PL APO CS2 63×/1.40 Oil objective at 0.75 base zoom with 1,024 × 1,024 pixels scan. Alexa488 and Atto647N-DOPE fluorescence were imaged simultaneously without any measurable bleed-through. Data were analysed using a custom ImageJ script that segments the Atto647N-DOPE channel to create a mask around the outer circumference of each bead. Segmented masks were then used to measure Alexa488 fluorescence around each bead.

### Cell culture

HeLa cells and MIA-Paca2 cells were cultured in DMEM supplemented with 10% FCS, and penicillin/streptomycin and passaged every 2-3days. NMuMG cells were grown in DMEM with 4.5 g/L glucose and 10 mcg/ml insulin and 10% FCS. For the generation of 3D spheroids, single cell suspensions were seeded in 8 well glass bottom imaging chamber (Ibidi) on a cushion of Matrigel (BD Bioscience) in 2% Matrigel containing growth media for 5days before being fixed and stained.

### Cell staining using recombinant biosensors

Cells were grown in 6well dishes on 24x24mm 1.5 glass coverslips to ∼80% confluence in full growth media. Media was removed and 4% PFA with 0.2% Glutaraldehyde (both EM grade) in PBS prewarmed to 37C was added for 20min at room temperature following by two quick washes and incubation with 50mM NH_4_Cl in PBS for 20min. After which the 6 well dish with coverslips were transferred onto ice and cells were washed with ice cold PIPES buffer. Cells were stained, blocked and permeabilised at the same time in 5% BSA + 0.5% (v/w) Saponin in PIPES with addition of labelled biosensors to a final concentration of 500nM for 45-60min on ice. For staining a standard metal heat block was inverted and cooled down in an ice bucket. A small piece of Parafilm was added onto the inverted heat block and 100ul drops of the staining solution were applied to the parafilm and placing the coverslips on top (with the cells facing the liquid). Afterwards the coverslips were transferred back into the 6well, on ice, and washed three times with ice cold PIPES. Cells were post fixed with 2% PFA in PBS on ice for 10min before being returned to room temperature for another 10min. Cells were washed 50mM NH_4_Cl in PBS at room temperature, rinsed with ddH_2_O, mounted, and sealed using Prolong Gold.

### Staining of Drosophila pupal wings

Wings were dissected from w[1118] (FlyBase: FBal0018186) pupae, aged for 28 hours after prepupae formation. These were fixed for 30 minutes in 4% paraformaldehyde/0.2% glutaraldehyde in PBS, the outer cuticle removed, and then quenched for 15 minutes in 50mM NH_4_Cl in PBS.

Subsequent staining steps were carried out on ice or at 4°C. Wings were blocked and permeabilised for 45 minutes in 5% BSA/0.5% Saponin in PIPES buffer. Recombinant biosensors were diluted in 0.1% Saponin in PIPES buffer (PI(4)P biosensor at final concentration of 150nM and PI(4,5)P_2_ biosensor at 500nM) with 1:200 Alexa 488 conjugated Phalloidin and incubated with the wings for 1 hour. Wings were then washed 12 times in 0.1% Saponin in PIPES buffer, post-fixed for 10 minutes in 2% PFA, washed a further 4 times with 0.1% Saponin in PIPES buffer, and mounted in 2.5% DABCO with 10% Glycerol in PBS.Wings were imaged on a Nikon A1R GaAsP confocal microscope using a 60x NA1.4 apochromatic lens within one week of mounting. Images were acquired posterior to longitudinal vein 4 with a pixel size of 110 nm and z sections spaced by 150 nm.

### Microscopy

Single colour staining using the recombinant biosensors conjugated with Alexa488 (Fig 3) was carried out using LSM880 Airyscan confocal microscope with a Plan Apochromat 63× 1.4 NA followed by standard 2D Airyscan processing. Multiplex staining using recombinant biosensors conjugated with Alexa488, 546 or 647 was carried out in full Z-Stacks of 150nm step size using LSM980 Airyscan with a Plan Apochromat 63x 1.4NA and processed using the joint deconvolution 3D Airyscan processing.

STED imaging was performed on a commercial expert line Abberior STED microscope equipped with 485, 561, and 645 nm excitation lasers and an Olympus Objective UPlanSApo 100× 1.40 NA. For two color STED of Arf1-Halo and PI(4)P, cells were seeded on a glass-bottom dish (3.5 cm diameter, No. 1.5 glass; Cellvis) coated with fibronectin (Sigma-Aldrich). Labelling with Halo substrate JF571 for was carried out for 1h using 1μM stocks. After the staining, cells were washed three times with growth medium to get rid of the excess dye and left for 1h in an incubator at 37°C and 5% CO_2_. Cells were then fixed and stained with biosensor against PI(4)P as described above. The 775 nm depletion laser was used to deplete dyes in two-color STED experiments. Two-color images were recorded sequentially line by line. Detection was carried out with avalanche photodiodes with detection windows set to 571–630 nm and 650–756 nm. Acquisition was carried out with Instruments Development Team, Imspector Image Acquisition & Analysis Software v16.3.16118 (http://www.imspector.de). The pixel size was set to 20 nm and Raw STED images were deconvolved to reduce noise using Richardson–Lucy deconvolution from the python microscopy PYME package (https://python-microscopy.org/).

### Electron microscopy

MIA-Paca2 cells were grown in media containing 10% FCS, treated with 1μM Apilimods in DMSO for 1h, fixed in 4%PFA with 0.2% Glutaraldehyde, pelleted and embedded in resin according to standard protocols. Pellets were stained with 1% osmium for 1h and 0.5% uranyl acetate overnight, embedded in epon resin, cut into 70-nm sections, and stained en bloc with uranyl acetate and lead citrate before imaging using an FEI Tecnai T12 Spirit at 80 kV.

## Acknowledgments

We acknowledge the Wolfson Light Microscopy Facility for the use of the LSM 980 Airyscan 2 supported by the Medical Research Council grant MR/X012077/1 and Christopher J Hill from the University of Sheffield Faculty of Science EM facility for support with this section electron microscopy. Hannes Maib is supported by a Wellcome Trust early Career award 225528/Z/22/Z. We further thank Prof. Jason King and Prof Elizabeth Smythe for helpful comments on the manuscript. RH is supported by BBSRC DTP studentship (BB/M011151/1), DS is supported by a Wellcome Trust Senior Fellowship (210630/Z/18/Z). D.H. Murray is supported by the Wellcome Trust grant 211193/Z/18/Z. P. A. is supported by the Forschungsgemeinschaft (German Research Foundation) grant SFB/TRR186 (Project A20).

## Notes

### Competing Interest Statement

The authors have declared no competing interest.

